# Resource limitation modulates the fate of dissimilated nitrogen in a dual-pathway Actinobacterium

**DOI:** 10.1101/364331

**Authors:** David C. Vuono, Robert W. Read, James Hemp, Benjamin W. Sullivan, John A. Arnone, Iva Neveux, Bob Blank, Carl Staub, Evan Loney, David Miceli, Mari Winkler, Romy Chakraborty, David A. Stahl, Joseph J. Grzymski

## Abstract

Respiratory ammonification and denitrification are two evolutionarily unrelated dissimilatory nitrogen (N) processes central to the global N cycle, the activity of which is thought to be controlled by carbon (C) to nitrate (NO_3_^-^) ratio. Here we find that *Intrasporangium calvum* C5, a novel menaquinone-based dual-pathway denitrifier/respiratory ammonifier, disproportionately utilizes ammonification rather than denitrification when grown under carbon or nitrate limitation, not C:NO_3_^-^ ratio. Instead, C:NO_3_^-^ ratio is a confounding variable for resource limitation. We find that the protein atomic composition for denitrification modules (NirK) are significantly cost minimized for C and N compared to ammonification modules (NrfA), indicating that resource limitation is a major selective pressure imprinted in the architecture of these proteins. The evolutionary precedent for these findings suggests ecological and biogeochemical importance as evidenced by higher growth rates when *I. calvum* grows predominantly using its ammonification pathway and by assimilating its end-product (ammonium) for growth under ammonium-deplete conditions. Genomic analysis of *I. calvum* further reveals a versatile ecophysiology to cope with nutrient stress and redox conditions. Metabolite and transcriptional profiles during growth indicate that transcript abundances encoding for its nitrite reducing enzyme modules, NrfAH and NirK, significantly increase in response to nitrite production. Mechanistically, our results suggest that pathway selection is driven by intracellular redox potential (redox poise), which may be lowered during resource limitation, thereby decreasing catalytic activity of upstream electron transport steps needed for denitrification enzymes. Our work advances our understanding of the biogeochemical flexibility of N-cycling organisms, pathway evolution, and ecological food-webs.

## Introduction

Globally, respiratory ammonification and denitrification are vital nitrogen (N) dissimilation pathways that either retain reactive N to support net primary productivity or close the N-cycle through the release of gaseous N, respectively [1]. The environmental controls on these two pathways, particularly the ratio of electron-donor to electron-acceptor (e.g., C:NO_3_^-^) [2], have gained attention [3–7] due to increased anthropogenic N inputs into the environment [8]. However, the effects of resource limitation on growth and pathway selection (i.e., allocation of C and N to dissimilatory and assimilatory processes), which are often confounded by C:NO_3_^-^ ratio, have not been tested. Strong selective pressures from Earth’s shifting biogeochemistry and oxidation-state have driven evolutionary adaptions to microbial electron transport chains (ETC) [9, 10], respiratory chain redox potentials [11–13], and protein atomic composition [14, 15], may shed light on how these pathways are regulated in contemporary organisms. Here, by identifying the biochemical and evolutionary differences between respiratory ammonification and denitrification, we disentangle the ecological significance and molecular mechanisms of electron transfer through either pathway in a dual pathway organism.

From a biochemical standpoint, the primary difference between respiratory ammonification and denitrification is their respective source of reducing equivalents in the ETC: 1) heme-based cytochrome c nitrite reductase used in respiratory ammonification receive electrons directly from the quinone (Q) pool [16] while 2) copper and *cd*_1_ nitrite reductases used in denitrification receive electrons from a soluble electron carrier (e.g., cytochrome c) via the bc_1_ complex [17]. From an evolutionary standpoint, we can place each N-module’s origin to a putative time in Earth history based on the metal co-factors that would have been bioavailable: heme-based cytochromes in an ancient, more reduced, environment compared to the copper-containing nitrite reductases in an oxidizing environment [18]. The bioenergetic chains of microorganisms also underwent selective pressure to shift from low-potential (LP) to high-potential (HP) quinones in response to Earth’s oxygenation [11, 12]. Menaquinone (MK) is thought to be the ancestral type of LP quinone [19]. Organisms that use ubiquinone (UQ) are thought to have evolved under high O_2_ tensions with *α-*, *β-*, *γ*-proteobacteria as the only bacterial clades to use UQ [12]. Surprisingly, our understanding for the biochemistry of denitrification is based predominantly on HP UQ-based systems [20], leaving a significant knowledge gap in the physiology and biochemistry of LP MK-based denitrifiers and how they link electron transfer with energy capture under resource limitation [21–23].

In order to resolve the mechanisms of C:NO_3_^-^ control on pathway selection and better understand branched respiratory chains in LP-based nitrate-reducing organisms, we undertook the characterization of the novel Gram-positive Actinobacterium strain *Intrasporangium calvum* C5: a dual-pathway nitrite reducer that uses MK as sole pool quinone. Here we show that over a range of C:NO_3_^-^ ratios, duplicated at two substrate concentrations, *I. calvum* disproportionately utilizes its ammonia-forming pathway during C limitation (≤ 0.4mM lactate), when C:NO_3_^-^ ratios are < 1 (an observation contrary to the current paradigm). Using a genome-guided approach coupled to time-series transcriptomics and metabolite profiles, we identified differentially expressed genes in the bacterium’s ETC and central metabolic pathways. Using this information to inform a metabolic reconstruction of the ETC and extensive literature on the biochemistry of the bc_1_ complex, we propose a new mechanism by which these two pathways are regulated at the biochemical level.

## Materials and Methods

### Culture Conditions

#### Media preparation

All cultures were grown at 30 °C and shaken at 250 rpm. Nitrate reducing minimal media was prepared with the following final concentrations: NaCl (0.6mM), NH_4_Cl (1.75mM) (for ammonium replete conditions but not used in NH4-deplete conditions), MgCl2 (0.2mM), CaCl2 (0.04mM), KCl (0.1mM), K2HPO4 (0.01mM), NaHCO3- (0.3mM), cysteine (1mM) as reducing agent, resazurin as redox indicator, and trace elements and trace vitamin solutions as reported [24, 25]. 1M sterile filtered (0.2μm) Concentrated stocks of 60% w/w sodium DL-lactate solution (Sigma-Aldrich, St. Louis, MO, USA), sodium-nitrate and sodium-nitrite (≥99%, Fisher Scientific, Pittsburg, PA, USA) were diluted into media prior to autoclaving to achieve the desired C:NO_3_^-^ ratio. C:NO_3_^-^ ratio was calculated based on [3] where the number of C atoms (n) in the e-donor is multiplied by the concentration of the e-donor, divided by the number of N atoms in the e-acceptor multiplied by the concentration of the e-acceptor (Table S4). See SI Materials and Methods for complete description of Hungate technique prepared media. Mean pH for all culture vessels (time series and end-point; Table S5), measured at the end of each experiment, was 7.3±0.05 (n=144).

### Analytical procedures

#### Growth Curve/Cell counts/Yield Measurements

Growth curves were measured from scratch-free Balch-tubes grown cultures using an automated optical density reader at OD_600_ nm (Lumenautix LLC, Reno, NV). End-point cultures were monitored until all replicates reached stationary phase (65-100 hours depending on C:NO_3_^-^ treatment) (Figure S6). Cell counts were performed by fixing cells in 4% paraformaldehyde (final concentration) for 20 minutes, filtered onto 0.2μm pore-sized black polycarbonate filters. A complete description is provided in SI Materials and Methods. Biomass concentrations were measured by filtration and drying as per standard protocol [26]. A complete description is provided in SI Materials and Methods.

#### Ion and Gas Chromatography Measurements

A dual channel Dionex ICS-5000+ (Thermo Scientific) ion chromatograph (IC) was used to measure organic (lactate, acetate, and formate) and inorganic (nitrite and nitrate) anions on an AS11-HC column and cations (ammonium) on a CS-16 column from the bacterial growth media. A complete description is provided in SI Materials and Methods.

### Phylogenetic, Genomic, and Transcriptomic Analysis

Genomic DNA was assembled using Canu (version 1.7.1) with an estimated genome size of 5 million base pairs [27]. The resulting single contiguous fragment was aligned to the *I.* calvum 7KIP genome (Acc: NC_014830.1) to compare sequence similarity in Mauve[28, 29]. Genome annotation for C5 was performed through the NCBI Prokaryotic Genome pipeline (www.ncbi.nlm.nih.gov/genome/annotation_prok/). Additional gene prediction analysis and functional annotation was performed by the DOE Joint Genome Institute (JGI) using the Isolate Genome Gene Calling method (Prodigal V2.6.3 February, 2016) under the submission ID 172966. The complete genome sequence and annotation is available in the NCBI database under the BioProject number PRJNA475609. A complete description of the phylogenetic, pathway analysis, and cost-minimization calculations is provided in SI Material and Methods. For transcriptomic analysis, the resulting raw reads were inspected using FastQC [30] to determine quality, read length, and ambiguous read percentage. Reads were trimmed based on quality score with a sliding window of 5 base pairs, quality cutoff of 28, trailing cutoff quality score of 10, as well as adapter contamination removal in Trimmomatic [31]. A complete description is provided in SI Materials and Methods. Statistical analyses were conducted in the R environment for statistical computing (r-project.org). Data that was tested using parametric statistical analysis were first validated for normality by visualizing the data as a histogram and testing via Shapiro-Wilks test for normality.

## Results

### Genomic analysis of *I. calvum* C5

We sequenced and analyzed the genome of *I. calvum* C5 to first compare its similarity to the type species *I. calvum* 7KIP. We identified a high degree of sequence similarity to 7KIP based on three homologous sequence regions as locally collinear blocks (SI Results). Genome size of C5 was 4,025,044 base pairs (bp), only 662 bp longer than 7KIP. Genomic analysis of the ETC revealed the typical suite of complexes common to facultative aerobes, including primary dehydrogenases (*nuo* complex, succinate dehydrogenase), alternative *NDH-2* NADH dehydrogenase, cytochrome bc_1_ complex, high-oxygen adapted cytochrome c oxidase (A-family), and low-oxygen adapted cytochrome *bd* oxidase. The bc_1_ complex subunits are also located immediately upstream of cytochrome c oxidase, suggesting that these enzymes are encoded in a single operon creating a supercomplex. Despite *I. calvum*’s seeming propensity for aerobic growth on a number of growth media [32], its bioenergetic system uses MK as its sole pool quinone. *I. calvum* also possesses multiple pathways for supplying electrons into the MK-pool, such as formate, malate, hydroxybutyrate, and glycerophosphate dehydrogenases. Once in the MK-pool, there are alternative pathways for MKH_2_ oxidation that can circumvent the bc_1_ complex, such as a membrane-bound respiratory nitrate reductase module (NarG). In addition, to NarG, its dissimilatory N module composition consists of a truncated denitrification pathway (N_2_O is a terminal product) using a copper nitrite reductase NirK and quinol-dependent nitric oxide reductase qNor. *I. calvum* also possesses both catalytic and membrane anchor subunits (NrfA and NrfH, respectively) for a pentaheme cytochrome c module involved in respiratory nitrite ammonification.

### *I. calvum* encodes for a functional NrfAH complex and assimilates NH_4_^+^ via respiratory nitrite ammonification

To gain insight into possible function of the NrfAH complex, we aligned the NrfA protein sequences from C5 and 7KIP to a collection of 33 recognized cytochrome c nitrite reductases from published annotated genomes (Table S1). This confirmed that NrfA from *I. calvum* is a member of the CxxCH 1^st^ heme motif group (Figure 1A), which forms one of four clades on the NrfA phylogenetic tree. We then queried the genomes of the taxa in our phylogeny for other annotated N-reducing modules used in nitrate reduction, nitrite reduction, NO-forming nitrite reduction, and primary pool quinone. Among the three major clades of NrfA, at least 5 additional taxa are noted having dissimilatory N-module inventories containing dual respiratory pathways: *S. thermophilum, B. azotoformans, B. bataviensis*, *B. bacteriovorus*, and *Candidatus* N. inopinata, (Figure 1A). None of the taxa in our NrfA phylogeny harbored the *cd*_1_ nitrite reductases (NirS). Due to the exclusive NirK representation in dual-pathway membership, we asked whether there might be differences in protein atomic composition between NirK and NrfA, given the disparate evolutionary origins of these modules [33]. We collected 20 additional publicly available NirK protein sequences from nondual-pathway denitrifiers (Table S1) and calculated the protein C and N composition for our NirK/NrfA collection as atoms per residue side-chain (Figure 1B). These results showed a significant depletion in C and N atoms per residue side-chain (ARSC) for NirK compared to NrfA (C and N: *p*<0.001; t-test), indicating that resource constraints are imprinted on the evolution of these proteins.

**Figure 1.**
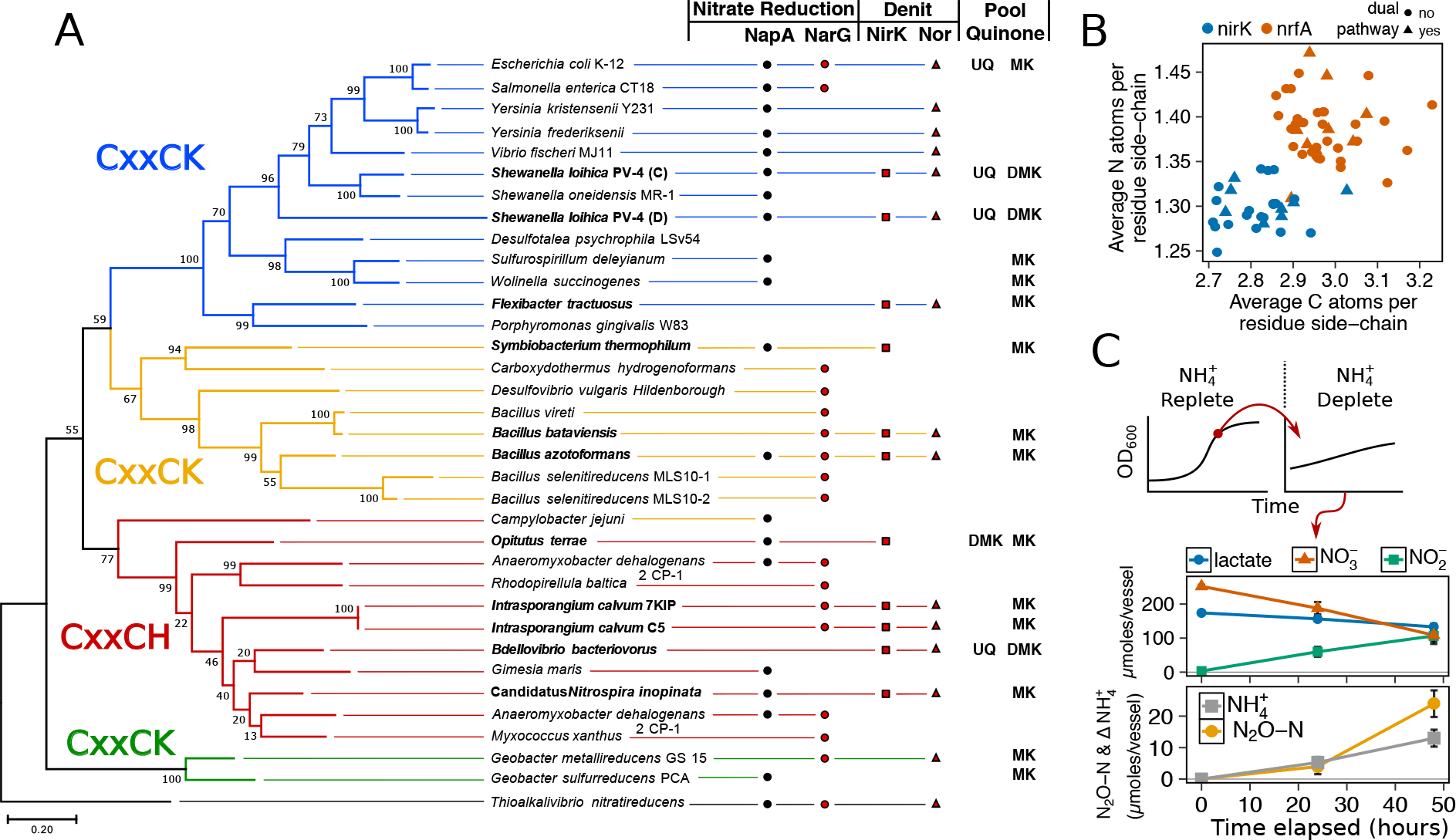
(A) Maximum likelihood phylogenetic tree of NrfA amino acid sequences from known respiratory ammonifiers and accompanying N-module composition for each organism. Pool quinone is also noted for dual-pathway nitrite reducers and model species. Colors of the main branches denote the 1^st^ heme motif type: CxxCK and CxxCH. (B) Protein atomic composition for N and C normalized to protein length for NirK and NrfA nitrite reductases. (C) State-transition from ammonium-replete to ammonium-deplete for *I. calvum* C5 grown under 8mM lactate 12mM nitrate minimal media at 30 °C. Metabolite profiles for ammonium-deplete are shown.

We next tested the functionality of *I. calvum*’s Nrf complex by growing the bacterium under reducing conditions (8 mM lactate, 12 mM nitrate, ammonium-replete). We then performed a state-transition where biomass from late-exponential growth phase was collected and anaerobically inoculated into ammonia-deplete media (Figure 1C; SI Results). Despite no detectable amounts of ammonium produced in the media over time, cell counts increased 5.4×10^5^±8.9×10^4^ cells/mL (0.126±0.02 optical absorbance at OD_600_) over a 48-hour incubation, indicating consumption of ammonium produced by NrfA. Net ammonium production was 13±2.7 μmoles with the remainder of dissimilated N being used by the denitrification pathway (24±4.2 μmoles N_2_O-N), resulting in a recovery of 97.4% dissimilated N. These results confirmed that *I. calvum* C5 has a functional Nrf complex and also consumes the product (ammonium) of respiratory nitrite ammonification.

### Respiratory nitrite ammonification exceeds denitrification under C-limitation

We investigated C:NO_3_^-^ control on respiratory ammonification versus denitrification on cultures of *I. calvum* C5 over a high resource C:NO_3_^-^ range (16-0.4 mM lactate, 12 mM nitrate; ratio 4-0.1) and low resource C:NO_3_^-^ range (1.6-0.04 mM lactate, 1.2 mM nitrate; ratio 4-0.1). This experimental design enabled us to evaluate C:NO_3_^-^ control over a broader range than previous studies that only considered ratios ≥ 1.5 [3, 4, 34], while also testing the effects of resource concentration on pathway selection. Under all the treatments tested, gas and ion chromatography measurements showed products of both respiratory pathways, differing only in the relative fraction of N_2_O versus ammonium production across treatments (Figure 2). At high resource concentrations, respiratory ammonification did not prevail at high C:NO_3_^-^ ratios (Figure 2A, Figure 2B, left panels; Table S2). Instead, significantly greater amounts of N_2_O were produced over ammonium, though nitrite was still the major extracellular end-product of nitrate respiration. Despite the predominance of N_2_O production under the high resource concentrations, ammonium production exceeded N_2_O production only at the lowest C:NO_3_^-^ ratio (0.4 mM lactate, ratio=0.1) (Figure 2) and accounted for 76.2±0.1% of dissimilated N.

**Figure 2.**
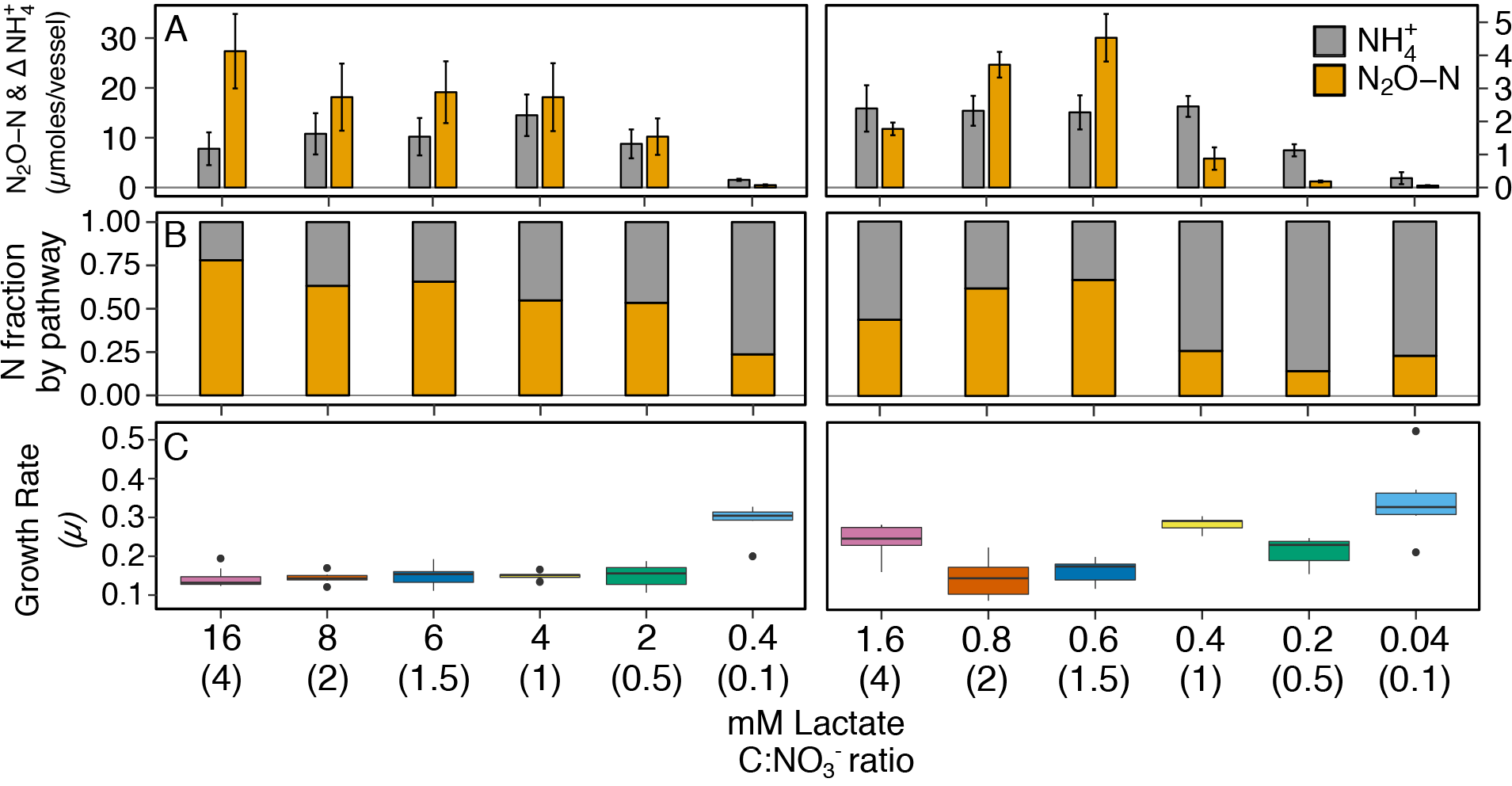
The effects of high resource (left; range of lactate concentrations with 12 mM NO_3_^-^) and low resource (right; range of lactate concentrations with 1.2 mM NO_3_^-^) concentrations with the same C:NO_3_^-^ ratio on pathway selection in *I. calvum* C5. (A) Production of N_2_O-N and net change of NH_4_^+^ over a 100-hour incubation period at 30 °C. Each bar represents the average of 8-10 replicates per treatment (Table S5). (B) Fraction of dissimilated N by pathway. (C) Growth rates for each corresponding treatment. The x-axis label defines lactate concentration and C:NO_3_^-^ ratio in parentheses.

Results from the low resource dataset provided weak support for the strict stoichiometry hypothesis that C:NO_3_^-^ controls pathway selection. Ammonia exceeded N_2_O production only under one high C:NO_3_^-^ ratio treatment (ratio=4; 1.6 mM lactate; Figure 2A, Figure 2B, right panels). However, at ratios ≤1 (≤0.4 mM lactate), significantly more ammonium than N_2_O was produced. On average, respiratory ammonification accounted for 78.1±8.9% of dissimilated N for lactate concentrations ≤0.4 mM. When these results are taken in context with cell physiology, we observed a significant and positive relationship between specific growth rate (*μ*) and the fraction of N dissimilated by respiratory ammonification (*R^2^*=0.5; *p*<0.001) (Figure 2C, S1; Table S3).

### Resource concentration influences the metabolite profiles of ammonium and N_2_O production

Given the co-occurrence of end products from both pathways during the end-point experiments (Figure 2), we next investigated the timing of ammonium and N_2_O production relative to metabolite profiles for lactate, nitrate/nitrite, and growth phase at two resource concentrations with the same ratio (8 mM and 0.8 mM lactate, ratio=2; Figure 1A, Figure S2, Figure S3). Despite ample e-donor and e-acceptor available for growth, the high resource cultures entered a quasi-stationary phase at ~50 hours, after which there was continued slow growth (Figure 1A). Metabolite profiles showed that ammonium and N_2_O production began simultaneously, as soon as nitrite was produced from nitrate reduction. The low resource cultures entered stationary phase at ~40 hours (Figure S2) after nitrate had been fully utilized. No further cell growth was observed after stationary phase was reached. These results show that cell growth occurred primarily on the reduction of nitrate, while nitrite reduction to ammonium and N_2_O occurred during a stationary growth phase, demonstrating that microbial activity is not always correlated with growth. The metabolite profiles for ammonium and N_2_O at low resources (Figure S2) did not mirror those observed at high resources (Figure 2A). The rate of N_2_O production significantly decreased and ammonium production oscillated rather than steadily increase through time. These differences in metabolite profiles, further demonstrate that concentration influences the activities of pathway bifurcation. Repeated time series experiments that were extended up to 300 hours show that nitrite is slowly depleted, but does not get fully consumed (Figure S3). When cultures were given nitrite, instead of nitrate as a terminal electron acceptor (8 mM lactate, 12 mM nitrite; ratio=2), we observed no immediate growth (as was observed with nitrate) but measured more N_2_O than ammonium production (33.4±4.8 μmoles N_2_O-N and 8.0±2.5 μmoles NH_4_^+^, respectively) (Figure S4), demonstrating respiratory ammonification does not exceed denitrification when nitrite is supplied as the sole acceptor in *I. calvum*.

### Nitrite-reducing modules are up-regulated during late exponential- and stationary-phase growth

In order to gain insight into mechanisms of gene regulation and transcriptional organization of *I. calvum*, we conducted RNA-Seq in parallel with the high resource time-series metabolite profile (Figure 3A). This approach enabled us to compare genome-wide differential expression based on log_2_ fold change (lfc) of RNA extracted from three growth phases: early exponential (EE), late exponential (LE), and stationary (ST) (Figure 3B, Figure S5). Within the central metabolic pathway beginning with the conversion of lactate to pyruvate, we observed a moderate decrease in transcript abundance of L-lactate dehydrogenase (LDH) (Intca_16740) between EE-LE and -ST (lfc = -1.6±0.7; -1.9±0.7), respectively. Lactate utilization protein C (LUP) (Intca_04080), an enzyme involved in lactate degradation, also showed a moderate and significant decrease in transcript abundance between EE-LE and -ST (lfc = -1.6±0.6; -2.4±0.6, *p*=0.002), respectively. *I. calvum* encodes for two parallel metabolic pathways for pyruvate conversion to acetyl-CoA: pyruvate dehydrogenase (PDH) (Intca_01255) and pyruvate ferredoxin oxidoreductase (PFOR) (Intca_15510). For PDH, there was a significant and moderate increase in transcript abundance between EE-LE and -ST (lfc = 2.1±0.6, *p*=0.002; 1.5±0.6), respectively. For PFOR, there was a minor decrease in transcript abundance between EE-LE (lfc = -0.43±0.5), and then a moderate increase in transcript abundance between EE-ST (1.1±0.5). Citrate synthase (Intca_04135), the enzyme catalyzing the conversion of acetyl-CoA to citrate and the first step of the tricarboxylic acid (TCA) cycle, showed a highly significant increase in transcript abundance between EE-LE and -ST (lfc = 4.3±0.5, *p*<0.001; 6.9±0.5, *p*<0.001).

**Figure 3.**
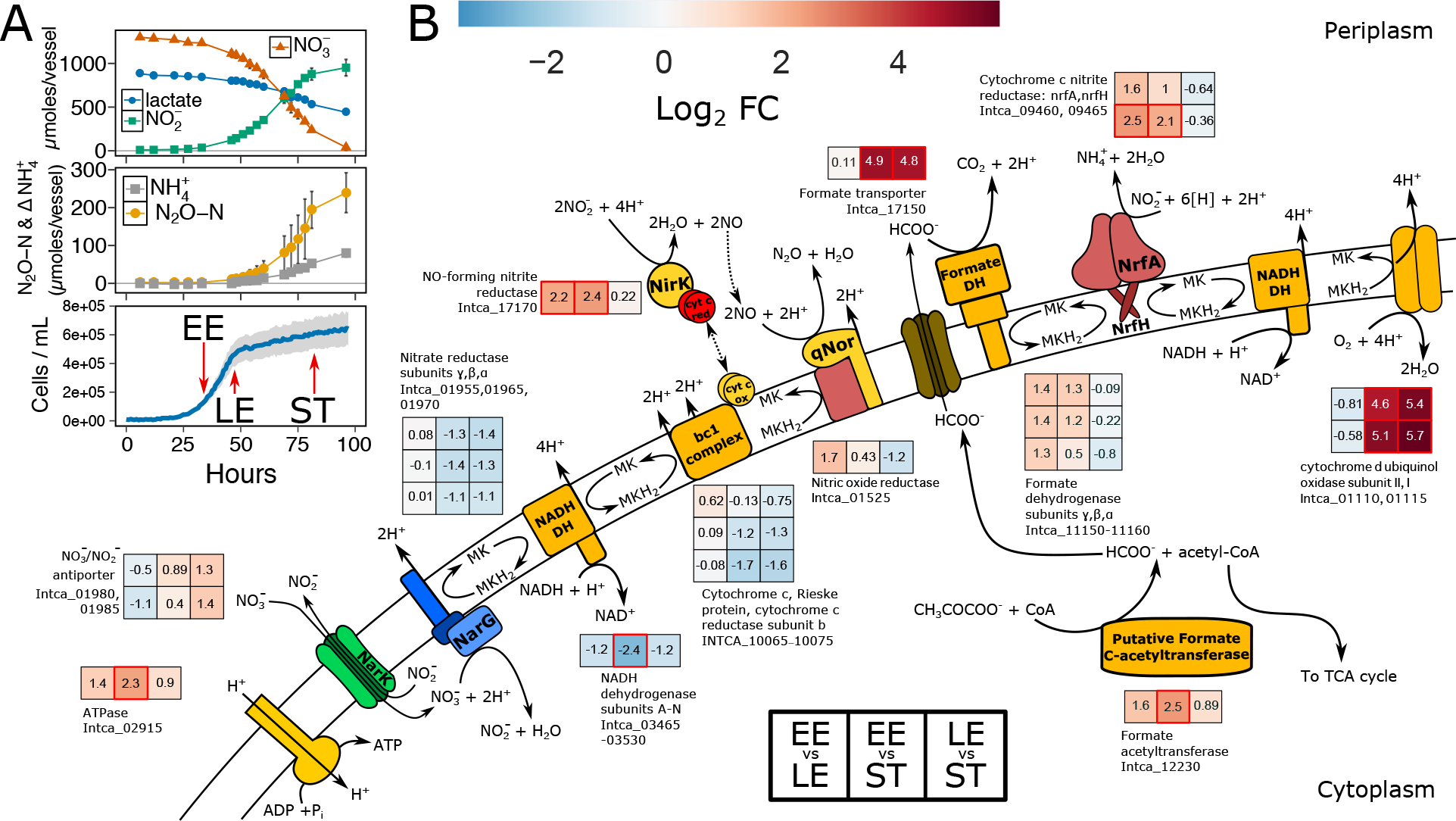
(A) Time-series metabolite profiles for lactate, nitrate, and nitrite (top pane), production of dissimilated end-products as N_2_O-N and net change in NH_4_^+^ ammonium production (middle pane), and corresponding growth curve of *I. calvum* cells grown under 8 mM lactate 12 mM nitrate (C:NO_3_^-^ ratio = 2) (bottom pane). Sampling points during growth phases are marked where transcriptomic profiling was performed (red arrows). (B) Metabolic reconstruction of the ETC from *I. calvum* with transcriptional changes for genes participating in dual-pathway dissimilatory nitrite reduction. Log_2_ fold changes in transcript abundance are shown for late exponential relative to early exponential growth phase (EE vs. LE), stationary phase relative to early exponential growth phase (EE vs. ST), and stationary phase relative to late exponential growth phase (LE vs. ST). Locus IDs for each gene product correspond to heat map subplots in the order shown (left-to-right for each growth phase and top-to-bottom for each locus ID specified). Higher transcript abundance is represented in red, lower transcript abundance in blue, and no change in transcript abundance in white. Significant changes in transcript abundance (*p* < 0.01) are marked as a red box. Value of log_2_ fold change is specified within each subplot. The log_2_ fold changes of 14 NADH dehydrogenase subunits (Intca_03465-03530) were averaged as transcriptional changes were all shifted in the same direction.

Within the ETC, there was moderate and significant decrease in transcript abundance for all subunits from the primary dehydrogenase (*nuo* complex; Intca_03465-03539) between EE-LE and -ST (lfc = -1.2±0.3; -2.4±0.6, *p*<0.001), respectively. Nitrate reductase subunits showed no change in transcript abundance between EE-LE (lfc = 0.01±0.07) and moderately decreased in abundance by ST (lfc = -1.2±0.1), which was corroborated by the depletion of nitrate during stationary phase. There was a significant increase in transcript abundance of *nirK* (Intca_17170) (lfc = 2.2±0.6, *p*=0.003; 2.4±0.6, *p*<0.001) and quinol dehydrogenase/membrane anchor subunit *nrfH* (Intca_09465) (lfc = 2.5±0.6, *p*=0.001; 2.1±0.6, *p*=0.003) by EE-LT and EE-ST, respectively, which coincided with nitrite production (Figure 3A). The catalytic subunit of the cytochrome c nitrite reductase complex (*nrfA*) (Intca_09460) also increased moderately in transcript abundance by EE-LT and EE-ST (lfc = 1.6±0.6; 1.0±0.6), respectively (Figure 3B). Contrary to the transcript abundance patterns of *nirK* and *nrfAH*, nitric oxide reductase (qNor; Intca_01525) transcripts moderately increased between EE-LT (lfc = 1.6±0.6) but decreased in the successive time periods (lfc = 0.43±0.6 between EE-ST; lfc = -1.2±0.6 between LE-ST) (Figure 3B).

There was a significant increase in transcript abundance of formate transporter *focA* (Intca_17150) between EE-ST, as well as LE-ST (lfc = 4.9±0.7, *p*=0.002; 4.8±0.7, *p*=0.002; respectively). We verified the production of formate in our ion chromatography measurements in the range of 100-200μM following late exponential growth. We also observed a moderate increase in transcript abundance of formate dehydrogenase (FDH) subunits (Intca_11150-11160). These results implicate the activity of formate oxidation, which would contribute to a ∆*p* in the periplasm via a Q-loop mechanism and the reduction of MK for electron transfer to nitrite via cytochrome c nitrite reductase. Considering that formate was not provided in our media recipe, an alternative pathway for formate production must exist in *I. calvum*. We also observed acetate production in similar concentrations as formate (100-200μM). In *E. coli*, formate is produced anaerobically from the action of pyruvate formate lyase (PFL). We identified a putative PFL based on genome annotation (Intca_12230), where transcript abundance also significantly increased by ST. PFL is also highly sensitive to oxygen [35], which was also in agreement with a significant increase in transcript abundance between EE-ST and LE-ST (Figure 3B) of cytochrome *bd* oxidase (Intca_01110 and Intca_01115), which is thought to protect anaerobic enzymes against oxidative stress [36].

## Discussion

We challenge the paradigm that C:NO_3_^-^ ratio controls pathway selection in a dual-pathway organism based on a simple principle: ratios do not account for the abundance of growth-limiting resources. We hypothesized that limitation in C or NO_3_^-^ should better predict pathway selection in a dual-pathway denitrifier/respiratory ammonifier. To test this hypothesis, we systematically measured the response of the Gram-positive Actinobacterium *Intrasporangium calvum* C5 to the same range of C:NO_3_^-^ ratios at both high and low resource loadings to better resolve mechanisms of pathway selection. We demonstrated that resource concentration, not C:NO_3_^-^ ratio, influences pathway selection. We found stronger support for respiratory ammonification preference under C-limitation (at low C:NO_3_^-^ ratios), which also grew at significantly higher growth rates (Figure 2). These results suggest that the NrfA complex, which receives electrons directly from the MK-pool, is optimized to maximize power when one or more resources are limiting. The enrichment of C and N ARSC in publically available NrfA over NirK protein seuqences (Figure 1B) provides further support and evolutionary precedence to ammonification preference over denitrification under resource limitation. This is because the end-product of ammonification can be used as an assimilatory N-source (Figure 1C), indicating no evolutionary constraint to cost minimize N. These data, together with metabolic reconstructions from metabolite and transcriptional profiles (Figure 3), suggest that C:NO_3_^-^ ratio alone is insufficient to explain pathway selection.

The theoretical basis for pathway selection is explained by the law of the minimum (LM) and the maximum power principle (MPP), which state that growth is limited by the least abundant resource and that biological systems are designed to maximize power in order to effectively allocate energy to reproduction and survival [37, 38], respectively. Here, it appears these two theories are working together: when resources are limited, the cell utilizes the respiratory pathway for growth that is optimized to maximize power. Power, in this case, is realized as higher growth rates from the cultures exhibiting disproportionately higher ammonium production than N_2_O production (Figure 2: high resources: C:NO_3_^-^ ratio = 0.1; low resources: C:NO_3_^-^ ratios = 4, 1, 0.5, 0.1). More specifically, the bacterium must generate a greater ∆*p* in order to maximize power when starved for a growth limiting resource. This may help to further explain how respiratory ammonification, which is overall energetically less favorable than denitrification (lactate with nitrite: ∆G° = -763.98 versus ∆G° = -1196.93, respectively), can have higher growth yields [39] and growth rates (Figure 2, Figure S1) under C- and N-limitation due to the higher energy yield on a per-nitrite basis (denitrification: -217 KJ per mole nitrite; respiratory ammonification: -399 KJ per mole nitrite). For comparison, a total of 8 H^+^ are translocated during denitrification by *I. calvum* (not including nitrate reduction since both pathways share this step) (Figure 3): NADH dehydrogenase translocates 4 H^+^ per MKH_2_ oxidized and the bc_1_ complex translocates an additional 4 H^+^ per MKH_2_ oxidized. However, 2 H^+^ must be consumed in the periplasm to reduce nitrite to NO [40]. qNor has a net zero H^+^ release (consumes 2 H^+^ to make N_2_O but releases 2 H^+^) without MKH_2_ regeneration [41]. Thus, a net total of 6 H^+^ are translocated per nitrite reduced in denitrification with added biosynthetic costs of making the bc_1_ complex and qNor. In respiratory ammonification, MK/MKH_2_ redox pair is cycled between NADH dehydrogenase and formate dehydrogenase. 6 electrons and 8 H^+^ are needed to reduce nitrite to ammonium, thus 3 MKH_2_ are needed [16]. If MKH_2_ is received from NADH dehydrogenase, 12 H^+^ are translocated plus 2 H^+^ from FDH. As each MKH_2_ is oxidized at the binding site of NrfH, 2 H^+^ are liberated [16], resulting in a net total of 12 H^+^ translocated per nitrite reduced for respiratory ammonification. This implies that the cell might deplete its NADH pool more rapidly on a per nitrite basis. However, if more protons are pumped in the early stages of growth, the cell would be allocating the ATP generated for anabolism, as evidenced by higher growth rates in the cultures exhibiting higher amounts of respiratory ammonification (Figure 2), which is supported by the MPP.

Under our high resource conditions (Figure 2; left panels), at C:NO_3_^-^ ratios ≥ 1, we observed that denitrification prevailed and these cultures had lower growth rates than the predominantly ammonium producing cultures. These high resource circumstances resulted in the production of toxic intermediates (i.e., NO_2_^-^ and possibly NO, albeit at undetectable levels), which may explain why these cultures had lower growth rates (Figure 2; left panels) and quasi-steady state growth curves in our high resource metabolite profile (Figure 3A). Rowley and colleagues [42] reported that at least 20% of the N_2_O released during high C conditions were produced by competition between nitrite and nitrate in the active-site of NarG. Under excess C concentrations, NarG produces intracellular NO from NO_2_^-^ and these intermediates are likely inhibitory to cell growth, which may explain why our growth curves (Figure 3A) reached a quasi-steady state before nitrate had been fully utilized (as compared to the low resource metabolite profile, Figure S2).

Furthermore, resources were not limiting growth under these conditions. Rather, the cells were likely experiencing toxicity from NO and NO_2_^-^ and thus the metabolic outcomes would be beyond the scope of the LM and MPP. Nonetheless, these results clearly demonstrate that end-product formation from the two resource concentrations tested, with the same C:NO_3_^-^ ratios, are not identical thereby refuting the C:NO_3_^-^ control hypothesis.

We selected a single treatment (8 mM lactate, 12 mM nitrate; C:NO_3_^-^ ratio = 2), in which we observed both denitrification and respiratory ammonification occurring simultaneously, for RNA-Seq in order to gain insight into the transcriptional organization of actively growing *I. calvum* cells (Figure 3). Strangely, we saw a decrease in transcript abundance encoding for two enzymes known to convert lactate to pyruvate, LDH and LUP. While normalized read counts (Figure S5) were generally consistent across growth phases, indicative of constitutive expression, further research investigating the mode of anaerobic lactate oxidation in *I. calvum* would illuminate how reducing equivalents are fed into its central metabolic pathway. For example, *S. loihica* PV-4 is known to use lactate for both denitrification and respiratory ammonification, but only uses acetate for denitrification [24]. Nonetheless, our transcriptomic data suggests that pyruvate plays a central role in providing reducing equivalents to the TCA cycle as Acetyl-CoA, as evidenced by significant upregulation in the genes encoding for pyruvate dehydrogenase and citrate synthase, as well as apparent “leaking” via incomplete lactate oxidation through the release of acetate and formate. Such leaking may be produced by a putative PFL, adding to the diversity of C utilization pathways feeding the ETC, and thereby driving pathway selection for nitrite reduction. Our transcriptomic results, coupled with a parallel metabolite profile (Figure 3), also suggest that the dual-pathway is induced by the presence of nitrite, and is not constitutively expressed like nitrate reductase, *narG*. Furthermore, it appears that the significant increase in transcript abundance for the gene encoding the *bd* oxidase helps to protect the anaerobic-dependent biochemical machinery against oxidative stress, thereby scavenging any residual oxygen during anaerobic growth.

Our metabolite profiles for N oxyanion respiration and N_2_O versus ammonium production show conflicting patterns relative to previous studies (Figure 3A, Figure S2). Yoon and colleagues [43] reported complete reduction of nitrate, production of nitrite, and then rapid consumption of nitrite, with N_2_O as the main end-product, by *S. loihica* PV-4 (5 mM lactate, 1 mM nitrate; ratio=0.6). When Yoon and colleagues [43] replaced nitrate with nitrite as the dominant electron acceptor (5 mM lactate, 1 mM nitrite, ratio=0.6), ammonification prevailed. Other research has shown the same response to nitrite replacement and ammonification dominance using non-fermentable C-sources (i.e., acetate) in chemostat enrichments of *Geobacter lovleyi*[44]. In our work, nitrite was never fully depleted (Figure 3A, Figure S2, Figure S3) and when nitrite was given as the only electron acceptor, the bacterium predominantly used denitrification but without concurrent growth (Figure S4). Similar to our work, Kraft and colleagues[34] also reported denitrification dominance when nitrite was supplied as the terminal acceptor. These differences highlight an incomplete understanding for the molecular mechanisms underlying the framework put forth by the LM and MPP.

A detailed look into the biochemistry of ETC complexes helps to shed light on the molecular mechanisms modulating pathway bifurcation. For example, Yoon and colleagues [3] demonstrated that elevated pH selects for ammonification in *S. loihica* PV-4. This phenotypic response is due to a decrease in the midpoint potential of the Rieske protein at higher pH [45–48]. Thus, any hindrance of electron flow through the bc_1_ complex would ultimately reduce the activity of downstream processes and promote alternative respiratory pathways. Nitrogen and C limitation have also been shown to influence flux distributions in redox sensitive proteins, including those found in electron transport [49]. A drop in the intracellular redox potential (redox poise) of the cell due to resource limitation may decrease the midpoint potential of the Rieske protein and reduce the activity of any downstream electron exit modules, such as NirK [50–52]. Thus, based on fundamental principles of protein redox chemistry and thermodynamics, it becomes clear that denitrification versus ammonification are likely not modulated by an arbitrary ratio of C:NO_3_^-^, but rather by thermodynamic constraints of the Q-cycle [11, 12]. The phenotypic response of higher rates of denitrification over ammonification at high C:NO_3_^-^ ratios in other published studies [3, 4] may also be due to enrichment bias for organisms that utilize quinones with higher midpoint potentials in their bioenergetic chains (Figure 1). Bergdoll and colleagues [11] suggested that comparisons of Rieske/cyt*b* complexes from organisms with high- and low-potential quinones may help to reconcile the thermodynamic properties of Q-cycle function. However, most of our understanding of denitrification bioenergetics is based on evolutionarily recent UQ-based HP bioenergetic chains from Gram-negative *α-*, *β-*, *γ*-proteobacteria. Because *I. calvum* uses a MK-based LP bioenergetic chain it may be possible that the differences in pathway selection across treatments are unique to LP chains.

Piecing together the evolutionary history of the N-cycle using isotopic signatures for geochemically available N module cofactors (i.e., Ni, Fe, and Mo) coupled to molecular evolutionary analysis has revealed respiratory ammonification was likely a major component of the Archean N-cycle [33]. Abiotic nitrite formation and depletion of ammonia through photodissociation [53] would have created selective pressures for a dissimilatory N pathway that also produced assimilatory N. We demonstrate that NrfA proteins are significantly enriched in N compared to NirK (i.e., no evolutionary constraints to cost minimize N in the *nrfA* gene product [15]) (Figure 1B) and that ammonium production (without accumulation in the medium) supports growth in *I. calvum* (Figure 1C). The Nrf module is also relatively simplistic in that it receives electrons directly from the quinol pool and not the bc_1_ complex used in denitrification. The early exit of electrons from the ETC (i.e., before reaching the bc_1_ complex) suggests that Nrf may have originated prior to the bc_1_ complex. Furthermore, the quinol oxidation site (Q_o_) of cytochrome *b* contains a PDWY motif, indicative of an ancestral LP respiratory chain found in many Gram-positive organisms [54]. However, there is still debate regarding the presence of a cytochrome *bc* complex in the last universal common ancestor [54, 55]. Lastly, the Nrf module is wired to operate via a q-loop with formate dehydrogenase whose Mo-cofactors would have also been bioavailable during the Archean, further supporting an early evolution.

In summary, we employ a new predictive framework that accounts for the biochemistry and evolutionary history of N modules, ETC complexes, and pool quinones to suggest the mechanisms by which these two pathways are regulated at the molecular level. With this understanding, it may be possible to extend our framework to environmental microbial populations and accelerate model development across different ecosystem scales (i.e., cross-scale systems biology).

## ACKNOWLEDGMENTS

We thank F. von Netzer, K. Hunt, S Morales, N. Stopnisek, K. Meinhardt, W. Qin, N Elliot, T. Hazen, H. Carlson, B. Ramsey, A. Murray, Z. Harrold, T. Morgan, and P. Longley for thoughtful feedback and discussions. This research was supported by a grant from the Nevada Governor’s Office of Economic Development (JG), by the Desert Research Institute (DRI) postdoctoral research fellowship program, and in part by Ecosystems and Networks Integrated with Genes and Molecular Assemblies (ENIGMA) (http://enigma.lbl.gov)—a Scientific Focus Area Program at Lawrence Berkeley National Laboratory under contract number DE-AC02-05CH11231 and funded in part by Oak Ridge National Laboratory under contract DE-AC05-00OR22725, and is based upon work supported by the U.S. Department of Energy, Office of Science, Office of Biological & Environmental Research.

## References

1. Knowles R. Denitrification. Microbiol Rev 1982; 46: 43–70.

2. Tiedje JM, Sexstone AJ, Myrold DD, Robinson JA. Denitrification: ecological niches, competition and survival. Antonie Van Leeuwenhoek 1982; 48: 569–583.

3. Yoon S, Cruz-García C, Sanford R, Ritalahti KM, Löffler FE. Denitrification versus respiratory ammonification: Environmental controls of two competing dissimilatory NO 3-/NO 2-reduction pathways in Shewanella loihica strain PV-4. ISME J 2015; 9: 1093–1104.

4. van den Berg EM, van Dongen U, Abbas B, van Loosdrecht MC. Enrichment of DNRA bacteria in a continuous culture. ISME J 2015; 9: 2153–2161.

5. Schmidt CS, Richardson DJ, Baggs EM. Constraining the conditions conducive to dissimilatory nitrate reduction to ammonium in temperate arable soils. Soil Biol Biochem 2011; 43: 1607–1611.

6. Hardison AK, Algar CK, Giblin AE, Rich JJ. Influence of organic carbon and nitrate loading on partitioning between dissimilatory nitrate reduction to ammonium (DNRA) and N2 production. Geochim Cosmochim Acta 2015; 164: 146–160.

7. Fazzolari É, Nicolardot B, Germon JC. Simultaneous effects of increasing levels of glucose and oxygen partial pressures on denitrification and dissimilatory nitrate reduction to ammonium in repacked soil cores. Eur J Soil Biol 1998; 34: 47–52.

8. Galloway JN, Aber JD, Erisman JW, Seitzinger SP, Howarth RW, Cowling EB, et al. The Nitrogen Cascade. Bioscience 2003; 53: 341.

9. Moparthi VK, Hägerhäll C. The evolution of respiratory chain complex i from a smaller last common ancestor consisting of 11 protein subunits. J Mol Evol 2011; 72: 484–497.

10. Dibrova D V., Cherepanov DA, Galperin MY, Skulachev VP, Mulkidjanian AY. Evolution of cytochrome bc complexes: From membrane-anchored dehydrogenases of ancient bacteria to triggers of apoptosis in vertebrates. Biochim Biophys Acta - Bioenerg 2013; 1827: 1407–1427.

11. Bergdoll L, Ten Brink F, Nitschke W, Picot D, Baymann F. From low- to high-potential bioenergetic chains: Thermodynamic constraints of Q-cycle function. Biochim Biophys Acta 2016; 1857: 1569–1579.

12. Schoepp-Cothenet B, Lieutaud C, Baymann F, Verméglio A, Friedrich T, Kramer DM, et al. Menaquinone as pool quinone in a purple bacterium. Proc Natl Acad Sci 2009; 106: 8549–54.

13. Soo RM, Hemp J, Parks DH, Fischer WW, Hugenholtz P. On the origins of oxygenic photosynthesis and aerobic respiration in Cyanobacteria. Science (80-) 2017; 355: 1436–1440.

14. Baudouin-Cornu P, Schuerer K, Marlière P, Thomas D. Intimate evolution of proteins: Proteome atomic content correlates with genome base composition. J Biol Chem 2004; 279: 5421–5428.

15. Grzymski JJ, Dussaq AM. The significance of nitrogen cost minimization in proteomes of marine microorganisms. ISME J 2012; 6: 71–80.

16. Einsle O, Messerschmidt A, Huber R, Kroneck PMH, Neese F. Mechanism of the six-electron reduction of nitrite to ammonia by cytochrome c nitrite reductase. J Am Chem Soc 2002; 124: 11737–11745.

17. Kraft B, Strous M, Tegetmeyer HE. Microbial nitrate respiration - Genes, enzymes and environmental distribution. J Biotechnol 2011; 155: 104–117.

18. Godfrey L V., Falkowski PG. The cycling and redox state of nitrogen in the Archaean ocean. Nat Geosci 2009; 2: 725–729.

19. Nitschke W, Dracheva S. Reaction center associated cytochromes in Anoxygenic Photosynthetic Bactera. In: Blankenship R, Madigan M, Bauer C (eds).1995. Kluwer Academic, Dordrecht, The Netherlands, pp 775–805.

20. Lalucat J, Bennasar A, Bosch R, Garcia-Valdes E, Palleroni NJ. Biology of Pseudomonas stutzeri. Microbiol Mol Biol Rev 2006; 70: 510–547.

21. Suharti, de Vries S. Membrane-bound denitrification in the Gram-positive bacterium Bacillus azotoformans. Biochem Soc Trans 2005; 33: 130–133.

22. Heylen K, Keltjens J, Stein LY. Redundancy and modularity in membrane-associated dissimilatory nitrate reduction in Bacillus. Front Microbiol 2012; 3: 1–27.

23. Decleyre H, Heylen K, Bjorn T, Willems A. Highly diverse nirK genes comprise two major clades that harbour ammonium-producing denitrifiers. BMC Genomics 2016; 17: 1–13.

24. Yoon S, Sanford RA, Löffler FE. Shewanella spp. Use acetate as an electron donor for denitrification but not ferric iron or fumarate reduction. Appl Environ Microbiol 2013; 79: 2818–2822.

25. Hillesland KL, Lim S, Flowers JJ, Turkarslan S, Pinel N, Zane GM, et al. Erosion of functional independence early in the evolution of a microbial mutualism. Proc Natl Acad Sci 2014; 111: 14822–14827.

26. APHA. Standard Methods for the Examination of Water and Wastewater, 23rd ed. 2012. American Public Health Association: Washington, DC.

27. Koren S, Walenz BP, Berlin K, Miller JR, Bergman NH, Phillippy AM. Canu: scalable and accurate long- - - read assembly via adaptive k - - - mer weighting and repeat separation. Genome Res 2017; 1–35.

28. Darling ACE, Mau B, Blattner FR, Perna NT. Mauve: Multiple Alignment of Conserved Genomic Sequence With Rearrangements. Genome Res 2004; 14: 1394–1403.

29. Darling AE, Mau B, Perna NT. Progressivemauve: Multiple genome alignment with gene gain, loss and rearrangement. PLoS One 2010; 5: e11147.

30. Andrews S. FastQC: A quality control tool for high throughput sequence data. Available online at: http://www.bioinformatics.babraham.ac.uk/projects/fastqc 2010.

31. Bolger AM, Lohse M, Usadel B. Trimmomatic: A flexible trimmer for Illumina sequence data. Bioinformatics 2014; 30: 2114–2120.

32. Kalakoutskii LV, Kirillova IP, Krassilnikov NA. a New Genus of the. J Gen Microbiol 1967; 48: 373-79–85.

33. Klotz MG, Stein LY. Nitrifier genomics and evolution of the nitrogen cycle. FEMS Microbiol Lett 2008; 278: 146–156.

34. Kraft B, Tegetmeyer HE, Sharma R, Klotz MG, Ferdelman TG, Hettich RL, et al. The environmental controls that govern the end product of bacterial nitrate respiration. Science (80-) 2014; 345: 676–679.

35. Abbe KS, Takahashi S, Yamada T. Invovlement of oxygen-sensitive pyruvate-formate lysase in mixed-acid fermetnation by Streptococcus mutans under strictly anerobic conditions. J Bacteriol 1982; 152: 175.

36. Das A, Silaghi-Dumitrescu R, Ljungdahl LG, Kurtz DM. Cytochrome bd oxidase, oxidative stress, and dioxygen tolerance of the strictly anaerobic bacterium Moorella thermoacetica. J Bacteriol 2005; 187: 2020–2029.

37. Lotka AJ. Contribution to the Energetics of Evolution. Proc Natl Acad Sci 1922; 8: 147–151.

38. DeLong JP. The maximum power principle predicts the outcomes of two-species competition experiments. Oikos 2008; 117: 1329–1336.

39. Strohm TO, Griffin B, Zumft WG, Schink B. Growth yields in bacterial denitrification and nitrate ammonification. Appl Environ Microbiol 2007; 73: 1420–1424.

40. Zumft WG. Cell biology and molecular basis of denitrification. Microbiol Mol Biol Rev 1997; 61: 533–616.

41. Hemp J, Gennis RB. Diversity of the Heme-Copper Superfamily in Archaea: Insights from Genomics and Structural Modeling. In: Schäfer G, Penefsky HS (eds). Results and Problems in Cell Differentiation. 2008. Springer Science+Business Media, pp 1–28.

42. Rowley G, Hensen D, Felgate H, Arkenberg A, Appia-ayme C, Prior K, et al. Resolving the contributions of the membrane-bound and periplasmic nitrate reductase systems to nitric oxide and nitrous oxide production in Salmonella enterica serovar Typhimurium. Biochem J 2012; 762: 755–762.

43. Yoon S, Sanford RA, Löffler FE. Nitrite control over dissimilatory nitrate/nitrite reduction pathways in Shewanella loihica strain PV-4. Appl Environ Microbiol 2015; 81: 3510–3517.

44. van den Berg EM, Rombouts JL, Kuenen JG, Kleerebezem R, van Loosdrecht MCM. Role of nitrite in the competition between denitrification and DNRA in a chemostat enrichment culture. AMB Express 2017; 7: 1–7.

45. Link TA, Hagen WR, Pierik AJ, Assmann C, von Jagow G. Determination of the redox properties of the Rieske cluster of bovine heart bc1 complex by direct electrochemistry of a water-soluble fragment. EurJBiochem 1992; 208: 685–691.

46. Ugulava NB, Crofts AR. CD-monitored redox titration of the Rieske Fe-S protein of Rhodobacter sphaeroides: pH dependence of the midpoint potential in isolated bc1 complex and in membranes. FEBS Lett 1998; 440: 409–413.

47. Zu Y, Couture MMJ, Kolling DRJ, Crofts AR, Eltis LD, Fee JA, et al. Reduction Potentials of Rieske Clusters: Importance of the Coupling between Oxidation State and Histidine Protonation State. Biochemistry 2003; 42: 12400–12408.

48. Trumpower BL. Cytochrome bc1 complexes of microorganisms. Microbiol Rev 1990; 54: 101–129.

49. Ansong C, Sadler NC, Hill EA, Lewis MP, Zink EM, Smith RD, et al. Protein redox dynamics during light-to-dark transitions in cyanobacteria and impacts due to nutrient limitation. Front Microbiol 2014; 5: 1–10.

50. Snyder CH, Merbitz-Zahradnik T, Link TA, Trumpower BL. Role of the Rieske Iron – Sulfur Protein Midpoint Potential in the Protonmotive Q-Cycle Mechanism of the Cytochrome bc 1 Complex. J Bioenerg Biomembr 1999; 31: 235–242.

51. Denke E, Merbitz-Zahradnik T, Hatzfeld OM, Snyder CH, Link TA, Trumpower BL. Alteration of the midpoint potential and catalytic activity of the Rieske iron-sulfur protein by changes of amino acids forming hydrogen bonds to the iron-sulfur cluster. J Biol Chem 1998; 273: 9085–9093.

52. Hunte C, Solmaz S, Palsdóttir H, Wenz T. A structural perspective on mechanism and function of the cytochrome bc1 complex. In: Schäfer G, Penefsky HS (eds). Results and Problems in Cell Differentiation. 2008. Springer Science+Business Media, pp 253–270.

53. Falkowski PG. Evolution of the nitrogen cycle and its influence on the biological sequestration of CO2 in the ocean. Nature 1997; 387: 272–275.

54. Kao WC, Hunte C. The molecular evolution of the Qo Motif. Genome Biol Evol 2014; 6: 1894–1910.

55. Dibrova D V., Cherepanov DA, Galperin MY, Skulachev VP, Mulkidjanian AY. Evolution of cytochrome bc complexes: From membrane-anchored dehydrogenases of ancient bacteria to triggers of apoptosis in vertebrates. Biochim Biophys Acta - Bioenerg 2013; 1827: 1407–1427.

56. White D. The physiology and biochemistry of prokaryotes, 3rd ed. 2000. Oxford University Press.

57. Thauer RK, Jungermann K, Decker K. Energy conservation in chemotrophic anaerobic bacteria. Bacteriol Rev 1977; 41: 100–180.

58. Giardina CP, Ryan MG. Total belowground carbon allocation in a fast-growing Eucalyptus plantation estimated using a carbon balance approach. Ecosystems 2002; 5: 487–499.

59. Edgar RC. MUSCLE: multiple sequence alignment with high accuracy and high throughput. Nucleic Acids Res 2004; 32: 1792–7.

60. Tamura K, Peterson D, Peterson N, Stecher G, Nei M, Kumar S. MEGA5: Molecular evolutionary genetics analysis using maximum likelihood, evolutionary distance, and maximum parsimony methods. Mol Biol Evol 2011; 28: 2731–2739.

61. Stamatakis A. RAxML version 8: A tool for phylogenetic analysis and post-analysis of large phylogenies. Bioinformatics 2014; 30: 1312–1313.

62. Kelley LA, Mezulis S, Yates C, Wass M, Sternberg M. The Phyre2 web portal for protein modelling, prediction, and analysis. Nat Protoc 2015; 10: 845–858.

63. Dobin A, Davis CA, Schlesinger F, Drenkow J, Zaleski C, Jha S, et al. STAR: Ultrafast universal RNA-seq aligner. Bioinformatics 2013; 29: 15–21.

64. Liao Y, Smyth GK, Shi W. FeatureCounts: An efficient general purpose program for assigning sequence reads to genomic features. Bioinformatics 2014; 30: 923–930.

65. Love MI, Huber W, Anders S. Moderated estimation of fold change and dispersion for RNA-seq data with DESeq2. Genome Biol 2014; 15: 1–21.

